# High frequency oscillations measured with optically pumped magnetometers from the human retina

**DOI:** 10.1101/2025.10.27.683907

**Authors:** James I. Lubell, Jens Voigt, Katharina Rolfs, Tilmann Sander, Sarang S. Dalal

## Abstract

Measurement of human neural activity with optically pumped magnetometers (OPMs) is rapidly proliferating, with sensitivity approaching that of cryogenic sensors. Most neuroscience research with OPMs to date has investigated neural responses below 70 Hz, but higher frequencies are also of interest. However, sensitivity to higher frequencies may be limited by both the inherent operating bandwidth of the current generation of zero-field OPMs as well as by their generally lower amplitude. To assess the upper bounds of OPM sensitivity we used the stereotypical retinal response to flashes of light. Retinal responses to light flashes characteristically exhibit a neural response above 70 Hz called the oscillatory potential (OP) when recorded with electroretinography (ERG). Here we adopt the term retinal high frequency oscillation (rHFO) to include measurement of similar activity with OPMs. Our comparison of magnetoretinography (MRG) and ERG shows that rHFO can be measured up to 140 Hz using rubidium-based zero-field OPMs.

## Introduction

Magnetoencephalography (MEG) systems based on optically pumped magnetometers (OPMs) have become commercially available and offer an attractive alternative to prevailing systems based on superconducting quantum interference devices (SQUIDs). While SQUIDs rely on liquid helium and associated cryogenic infrastructure, OPMs operate at room temperature and, much like electroencephalography (EEG) electrodes, are small and light enough to be placed as needed anywhere on the body (Barratt et al., 2018). Their form factor furthermore allows OPMs to measure from locations that are inaccessible to most modern MEG SQUID systems. (Knappe et al., 2014; Westner et al., 2021, Marquetand et al., 2021). While in many aspects OPMs can be used in place of SQUIDs, the bandwidth and sensitivity of OPMs is limited by their underlying physics, including the magnetic moment of the atoms used, the number of atoms present in the measurement, and the speed and frequency of how those particles return to equilibrium (Budker & Romalis, 2007). Recent work has shown that OPMs operating in closed-loop modes can yield higher signal to noise ratios (SNR) across a wider bandwidth, but the relative measurement bandwidth of OPMs depends largely on the background fields it is operating in (Iivanainen et al., 2020).

In standard operation increasing an OPM sensor’s sensitivity effectively decreases bandwidth and vice versa (Iivanainen et al., 2020). OPMs that are commercially available for MEG specify slightly stronger intrinsic noise than SQUIDs (Marhl et al. 2022). Of the major commercial OPM manufacturers, QuSpin states that their Gen-3 sensors have a sensitivity of 7–10 fT/√Hz from 3-100 Hz (dual-axis mode, https://quspin.com/products-qzfm/) and FieldLine reports that their sensors have a sensitivity of ∼15 fT/√Hz, from DC to 150 Hz (Robinson et al., 2022). In addition to these specifications, the measurement of low-amplitude neural signals also depends on the extent of environmental interference as well as susceptibility to motion artifacts from any remnant gradients that remain uncorrected in the shielded room. (Iivanainen et al., 2020; Robinson et al., 2022).

Iivanainen et al. (2020) reported that OPM-MEG over the occipital cortex detected visual task-induced activity in the 40–70 Hz range with greater amplitudes than SQUID-MEG. This included enhanced power differentiation in the 7–13 Hz and 70–130 Hz bands at both the source and sensor levels and estimated that maintaining sensor noise at 3 fT/√Hz within a ∼10 nT background field in a three-layer magnetically shielded room (MSR) was adequate for reliable detection of high-gamma activity (60-70 Hz, Iivanainen et al., 2020). In lieu of improving shielding, more recent studies have demonstrated that optimizing OPM sensor placement and increasing channel density can enhance SNR and can achieve broadband gamma detection (up to about 80 Hz) comparable to conventional MEG (Hill et al. 2024).

These findings indicate that improved control over noise enables higher SNR measurements of high frequency activity; however, the maximal bandwidth of 130 Hz and the identifiable neural signals at approximately 80 Hz, do not establish an upper frequency limit for OPM measurements.

Closed-loop OPM measurements, a feature available in all commercial OPM-MEG systems, indirectly offers another improvement on SNR through ongoing dynamic background field compensation (Robinson et al., 2022, Seymour et al., 2022). In standard open-loop operation, OPM sensors are less robust to fluctuations in the background field, which result in nonlinear magnetometer responses that may negatively affect SNR. In closed-loop mode an acceptable SNR is still achievable despite strong background gradients (Iivanainen et al., 2019) or changes in the residual background field over time. This is implemented by dynamically adjusting current fed to the nulling coils embedded in each sensor package such that the field impinging on the magnetometer remains zero despite environmental fluctuations (Alem 2013, Fourcault 2021, Robinson 2022).

The stated bandwidth for sensors used in this study in open-loop mode is DC to 150 Hz (FieldLine Inc., Boulder, Colorado, USA). Activating closed-loop mode can increase the nominal bandwidth of these sensors to 350 Hz. The usable bandwidth in closed-loop mode is influenced by the gradients present and the gain coefficients that optimize the flatness of the sensor’s frequency response (Alem et al., 2023). While closed-loop mode nominally increases bandwidth to DC - 350 Hz, background fields may lead to noise magnification above 150 Hz. This magnification reduces the closed-loop frequency response to a range similar to open-loop mode (Alem et al., 2023). Crucially, in comparison to open-loop mode, closed-loop mode can maintain consistent SNR across the available frequency response range (Robinson et al., 2022).

The ability to measure reliably above 70 Hz in more standard shielded environments (i.e., 3 layer MSRs) makes OPMs a viable method to measure epileptic spikes (Feys et al., 2023), visual attention (Anders et al., 2025), and peripheral nervous system recordings (Bu et al., 2022), with potential for e.g. auditory brainstem responses and high-frequency retinal oscillations. (For a review of high frequency brain activity in M/EEG see Muthukumaraswamy, 2013 and see Brickwedde et al., 2024 for a perspective on clinical applications of OPMs and Wang et al., 2024 for a comparison of visual and auditory responses with SQUIDs and OPMs.)

Electroretinography (ERG) is a clinical technique that measures the summed neural responses of the retina and can be a useful diagnostic of retinal health and retinal processing (Robson et al., 2018). A major drawback of conventional ERG is the comfort of the electrodes; in clinical settings usually either a fiber electrode placed on the sclera (Dawson Trick Lawson electrode, DTL), or a Burian-Allen rigid contact lens electrode is used (Robson et al., 2022). Both DTL and Burian-Allen electrodes require direct contact with the eye and some associated discomfort (Robson et al., 2018); typically Burian-Allen electrodes are used together with local anesthesia. Promisingly, our lab has shown that the prominent retinal evoked responses to light flashes typically measured with ERG, the a- and b-waves can also be measured with an OPM-based magnetoretinogram (MRG) without even coming into direct contact with the participant (Westner et al., 2021). The first inflection, the a-wave, peaks about 25 ms after light flash onset and is thought to arise from the photoreceptor response (Inui et al., 2006). This is followed by the b-wave potential which is thought to arise from the ON bipolar cells, typically 50-80 ms after light onset (Frishman, 2013; Luo & Frishman, 2011; Robson et al., 2022). These measurements were achieved in a 3-layer magnetically shielded room (Vacuumschmelze Ak3b) and showed that oMRG (OPM-based MRG) was comparable to ERG for lower frequency signals (Westner et al., 2021).

In this study, we aimed to measure the magnetic counterpart of the high-frequency ERG waveform known as the oscillatory potential (OP). The OP is an oscillatory burst that peaks around 120 Hz in ERG recordings and rides along the rising positive b-wave inflection (Wachtmeister, 1998). The OP typically begins at approximately 19-20 ms after light onset and ends around 45-50 ms (Robson et al., 2022). It is clinically relevant in diseases that do not symptomatically impact the a- and b-wave components of the retinal response. For example, patients with diabetic retinopathy, the world’s leading cause of blindness, exhibit alterations in the OP that can be the sole indicator of disease progression (McAnany et al., 2019). Other diseases in which a degraded OP is an early indicator include congenital stationary night blindness and retinopathy of prematurity (Wachtmeister 1998; Fulton et al., 1987). It would also facilitate retinal measurements in patients with neurodegenerative diseases (Alves et al., 2024; Soto Linan et al., 2025) and psychiatric disorders (Hébert et al., 2010). Consequently, oMRG’s ability to measure high frequency components would be highly desirable for potential use in clinical diagnostics as well as research. In addition, it would demonstrate the quality with which OPMs can measure high-frequency neural activity in general, with relevance to cortical OPM-MEG measurements.

Since *oscillatory potential* would be a misnomer when measured with magnetometers, and since the terminology is unspecific to the retina in the wider MEG community, we have chosen to use the term *retinal high frequency oscillations* (rHFO) when referring to either the ERG-measured OP or its OPM-measured magnetic counterpart. Exactly which retinal neurons contribute to the rHFO is debated, with the two prominent views arguing either in favor of retinal ganglion cell sources (Yokoyama et al., 1999; Kenyon et al., 2003) or from the direct and indirect contributions of bipolar, amacrine, and interplexiform cells (Wachtmeister, 1998). Regardless of its origin, the rHFO are as robust as the a-b-wave response to a flash of light and can also be measured without directly contacting the eye (Heinrich & Bach, 2004; Wachtmeister, 1998; Westner et al., 2021). That rHFO characteristically is within the 75-160 Hz range and peaks at 120 Hz at around 50 ms after light flash onset (Frishman 2013, Gauthier et al., 2019), making it a suitable candidate for testing the claimed operational OPM bandwidth of DC - 350 Hz in closed-loop mode (Alem et al., 2023; Knappe et al., 2019, Robinson et al., 2022). In our previous MRG study rHFO results were difficult to verify (Westner et al. 2021), but we hypothesized that an environment free of strong background gradients and with less environmental noise would enable OPMs to measure the rHFO.

To that end, we measured retinal responses in the Berlin Magnetically Shielded Room Two (BMSR-2), which has remnant fields below 1 nT (Bork, 2001). The BMSR-2 has seven layers of µ-metal that provide, at frequencies less than 0.01 Hz, a passive shielding factor of 75,000. In addition, the active shielding surrounding the measurement chamber dampens magnetic interference by a factor of more than a million (Altarev et al., 2014). The ideal environment of the BMSR-2 and the use of closed-loop mode suggested that high-frequency physiological signals would have sufficient SNR to be measured with FieldLine OPMs, possibly up to 350 Hz.

The novelty and growing use of OPMs for physiological measurements necessitates quantifying what is currently possible and what reasonable targets for examination are, given the limits of this technology. In this paper, we show that MRG measurements using OPMs indeed capture high frequency signals and suggest a practical upper sensitivity limit for currently available OPMs.

## Methods

### Participants and measurement setup

Out of the ten participants recruited, one participant was excluded after OPMs were not initialized properly prior to recording, thus leading to exceptionally noisy data. The remaining participants (2 female; mean age 38.2 years, SD 12.9) had normal or correctable-to-normal eyesight. Participants did not wear glasses (to prevent magnetic interference) or contact lenses (for compatibility with ERG measurements); this was not expected to influence retinal responses, as stimuli were full-field light flashes that contained no visual detail.

Measurements were performed in the Berlin Magnetically Shielded Room Two (BMSR-2), located at the Physikalisch Technische Bundesanstalt (PTB) in Berlin, Germany. A total of 14 OPM sensors (V2 sensors; FieldLine Inc., Boulder, Colorado, USA; dimensions of one sensor: 30 x 15 x 12 mm) were placed in a 3D-printed holder shaped to surround the eye (see Figure 1). The OPM holder was then affixed to the the participants’ head with a band such that OPM 9 was positioned as best as possible over the left outer-canthus, OPM 5 along the midline axis of the eye on the eyebrow, and OPM 12 directly below the eye on the same midline axis (Figure 1). A small amount of variation relative to these landmarks resulted from individual differences in head shape. Sensor location choice was guided by Westner et al. (2021) and Katila et al. (1981).

**Figure 1.**
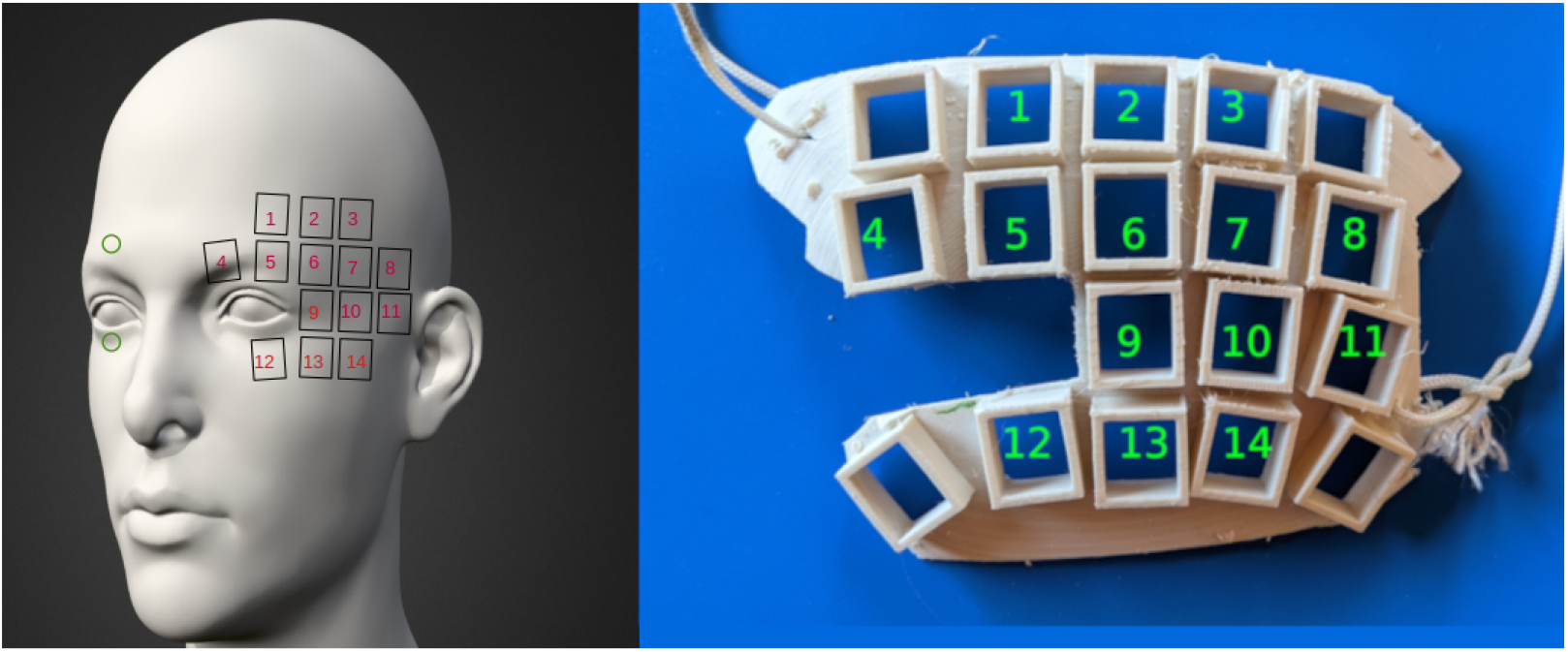
Sensor placement. **Leftside**: ERG and MRG layout and numbering. Above eye and on eyelid ERG electrodes are represented with green circles (outer right canthus ERG reference electrode not depicted). OPM locations and numbering are represented by black squares (empty and non-functioning sensors in the array are not depicted). Each participant rested their chin to help stabilize head and reduce movement. **Rightside**: Print of MRG array used showing empty slots.

ERG and EOG were measured from the right eye using 5 mm silver cup electrodes (Spes Medica, Genova, Italy). An electrode for measuring ERG was attached to the participants’ lower right eyelid. Although clinical ERG commonly employs an on-eye electrode (contacting the cornea or sclera), we opted for a lower eyelid-based ERG to maximize participant comfort and since we expected minimal environmental noise also in electrode measurements due to the particularly robust shielding of the BMSR-2 (Coupland & Janaky, 1989; Esakowitz et al., 1993; Kakisu et al., 1986; McCulloch et al., 1997; Porciatti & Ventura, 2004). Another electrode was attached directly above the right eyebrow to capture the vertical electrooculogram, a reference electrode was placed at the right outer canthus, and a ground electrode was attached to the inner right forearm close to the wrist. Electrodes were connected to a custom-built battery-driven EEG amplifier (-3 dB bandwidth 0 to 5 kHz, 0.16 Hz HP cut-off filter, AD8221 input chip, 24 bit ADC) and activity measured by the OPMs was recorded with the FieldLine system. The EEG and OPM systems each used a sampling rate of 1000 Hz, with common stimulation triggers used to synchronize data from each system offline.

Each participant was seated in the BMSR-2 and placed their chin on a board 85 cm from the wall onto which the flash stimuli was presented. While the electrodes were attached prior to entry into the BMSR-2, the OPM sensors mounted in their array were placed on the participant and secured inside the BMSR-2. After closing the BMSR-2 doors, background lighting was dimmed, and the OPM sensors were recalibrated for zero field. Following a successful fine zeroing, the participant was told the experiment was about to begin, the lights were turned off, the EEG and OPM recordings were started, and a light flash sequence was initiated.

### Stimuli

The BMSR-2 is equipped with four high performance ColdVision Light Source (CV-LS) units (Schott AG, Mainz, Germany). Each CV-LS is a high intensity fiber-optic LED light source that can be precisely controlled via a digital interface. Each light source was connected to fiber optic illuminators, originally installed for general purpose lighting within the BMSR-2. We took advantage of two of the illuminators to deliver light flash stimuli to the participants. We produced full-field flash stimuli by aiming the heads of two of the illuminators at a single spot (84.8 cm away) on the wall and programmatically controlling the corresponding CV-LS units. The output intensity of the stimulating CV-LS units were lowered to 70% with a pulse duration of 694 µs, which resulted in a flash intensity of 0.72 cd·s/m^2^.

Stimulus timing and triggering were controlled with an Arduino Nano (ATmega328, AVR architecture, 16 MHz clock speed). The code flashed to the Arduino Nano used analog pins to simultaneously turn two of the CV-LS units on (through the rear analog port), send out a 5-volt trigger pulse with an up time of 5.0 ms, and then turn off the CV-LS units after 694 µs. The trigger output was split and sent to both the EEG amplifier and OPM system. The period that the light was off was randomly jittered between 1.0 s and 1.1 s. A total of 650 trials were allowed to elapse before the Arduino Nano was turned off. Trial progress was monitored in the Arduino serial monitor.

### Analysis

Analysis was done using the MNE Python toolbox (MNE-Python Developers, 2022), MATLAB, and custom scripts.

### Preprocessing - Artifact detection

Initial preprocessing was performed prior to bandpass filtering for the HFO, in order to catch artifacts and anomalous epochs that are less evident in the HFO band. First, both the EEG and OPM datasets, which were recorded separately, were epoched (-200 to 300 ms) around the trigger issued by the Arduino that was concurrent with light onset. Second, the EEG and OPM datasets were merged, highpass filtered at 2 Hz, and visually inspected for artifacts. Finally, all trials with blinks, eye movements, muscle twitches and other artifacts were flagged for later removal. An average of 587.2 trials (SD 34.9) per participant remained after visual artifact rejection.

### Evoked responses

#### High frequency activity

Continuous data were bandpass filtered from 110 – 145 Hz with a one-pass, zero-phase, non-causal bandpass FIR filter, Hamming window 0.0194 passband ripple with 53 dB stopband attenuation, a lower transition bandwidth of 27.50 Hz (-6 dB cutoff frequency: 96.25 Hz), an upper transition bandwidth of 36.25 Hz (-6 dB cutoff frequency: 163.12 Hz), and filter length of 121 samples (121 ms). After filtering, continuous data were epoched and the EEG and OPM datasets were merged. The artifactual epochs identified during preprocessing were now rejected.

#### Low frequency activity

The same process was performed again, but this time data were filtered from 1 to 40 Hz with a one-pass, zero-phase, non-causal bandpass FIR filter, Hamming window 0.0194 passband ripple with 53 dB stopband attenuation, a lower transition bandwidth 1.00 Hz (-6 dB cutoff frequency: 0.50 Hz), an upper transition bandwidth of 10.00 Hz (-6 dB cutoff frequency: 45.00 Hz), and a filter length of 3301 samples (3.301 s). Filtering prior to epoching avoided introducing unwanted edge artifacts (Gross et al., 2013).

### Inter-trial Phase Coherence (ITPC)

#### ITPC rHFO

A time-frequency representation of the unfiltered and epoched data was computed using Morlet decomposition (mne.time_frequency.tfr_morlet). Ten logarithmically spaced frequencies between 110 and 145 Hz were generated using the base 10 logarithm ([110, 113.4, 117, 120.6, 124.4, 128.2, 132.2, 136.4, 140.6, 145]). The number of cycles was varied for each band by dividing each frequency by nine (cycles per frequency: [12.2, 12.6, 13, 13.4, 13.8, 14.2, 14.7, 15.1, 15.6, 16.1]). Power and ITPC were inspected for each participant. To evaluate whether oscillatory activity was consistent across subjects, a group ITPC measure was derived. The ITPC of each participant was Fisher-transformed prior to averaging across the participants, the resulting grand average was then inverse transformed back into ITPC. Using a similar process and to ensure completeness, ITPC was assessed from 110-170 Hz, and 110-300 Hz 15 cycles.

An additional ITPC analysis was performed to characterize the frequency response of the OPMs. Continuous data were highpass filtered at 1 Hz and epoched using the identical parameters as the above (See preprocessing). For each subject at each sensor the amplitude spectrum of the evoked response (5 to 100 ms post-stimulus) was computed using a Hanning-windowed Fast Fourier Transform. For each subject at each sensor the same procedure was followed using the baseline period (−50 to −5 ms). The standard error of both spectra was computed as the standard deviation of each spectrum divided by the square root of the number of participants.

#### ITPC Statistics

Using a Mann–Whitney U test, the significance of ITPC results was calculated. For each individual, an initial U-test compared each ITPC time point per channel per band to its relative baseline period (-50 - 0 ms). The resulting statistical values were then used for the group statistical test. For the group test another U-test was used to compare the same time point from all participants to the pooled statistical values of the baseline period across all participants. With 10 participants and a 50-sample baseline period, the group statistical test compared 10 samples for any given latency to the 500 pooled baseline period statistical points. Because the polarity of the signal is not preserved when computing the individual Mann–Whitney U test or in the ITPC data, and we are interested in examining changes from baseline, it was necessary to reintroduce a measure of polarity prior to computing the group statistic. This was achieved by taking the median value of a participant’s ITPC baseline period in each band and channel and determining if it was less than or equal to any time point in that same band and channel. If a time point was less than or equal to the median baseline period it was multiplied by negative one, thus reintroducing the polarity of the signal prior to it being entered into the group statistic. Following this process the results were then corrected for multiple comparisons using the False Discovery Rate (Benjamini & Hochberg, 1995).

### Mutual Information (MI)

To assess the amount of information shared between the under-eye ERG electrode and the OPM sensors, mutual information was computed both at the participant and group level. Three perspectives of the MI findings are provided: MI at the participant level using the Hilbert envelope of the data band-pass filtered from 130-145 Hz (see filter settings above), MI at the group level, and MI at the group level using the phase data that was extracted for the ITPC.

#### MI per participant

Mutual information was then computed between the under-eye ERG electrode and each sensor, including the over-eye electrode, from -50 to +120 ms on a per participant level (Cohen, 2014). The number of bins per participant was estimated with the Freedman-Diaconis rule and showed that the average number of bins was 8.64 (median 9, SD 1.59, range 6 - 14); rounding up, this determined the use of 9 bins for all per-participant mutual information computations.

#### MI Group

To compute the mutual information across all participants the grand average was computed across the evoked Hilbert envelope data, mutual information was again computed between the under-eye electrode and each sensor, including the over-eye electrode, from -50 to +120 ms. The number of bins was estimated with the Freedman Diaconis rule and showed that the average number of bins for each sensor was 9.6 (median 9, SD 1.24, range 8 - 12), thereby determining 10 bins.

#### MI group phase

To assess the MI between the under-eye electrode and the other sensors, phase data was first extracted per participant following the same process as the ITPC computations. For each participant the under-eye ERG data was compared to every other sensor by sliding a 25 ms window in 1 ms steps over the data within each frequency band and channel. The resulting findings, as expressed in bits, were then averaged together to deliver the group average (Frequency Bands: [110, 113.4, 117, 120.6, 124.4, 128.2, 132.2, 136.4, 140.6, 145]).

#### Mutual Information Statistics

To determine statistically significant shared information at both the participant and group level comparisons, a permutation test was used. On each permutation the data being compared was randomly shifted around to create a new distribution and a new MI comparison was computed. After 5000 repetitions the permuted MI information created a null distribution. A z-score was then derived by subtracting the mean of the permuted MI from the MI and dividing that number by the standard deviation of the permuted MI.

## Results

The retinal a-wave and b-wave were clearly observed in the OPM measurements, with the oMRG resembling the ERG across participants (Figure 2). Out of the sixteen OPMs, the sensor with the highest b-wave amplitude per participant was included in the average and is plotted with the ERG electrode. In the group average, the a-wave amplitude peaked at 18 ms in the oMRG and peaked at 17 ms in the ERG. The b-wave peaked at 79 ms, as measured by both oMRG and ERG (Figure 2). The correspondence between oMRG and ERG replicates our earlier report (Westner et al. 2021) in which we show that OPMs can measure retinal low frequency signals.

**Figure 2.**
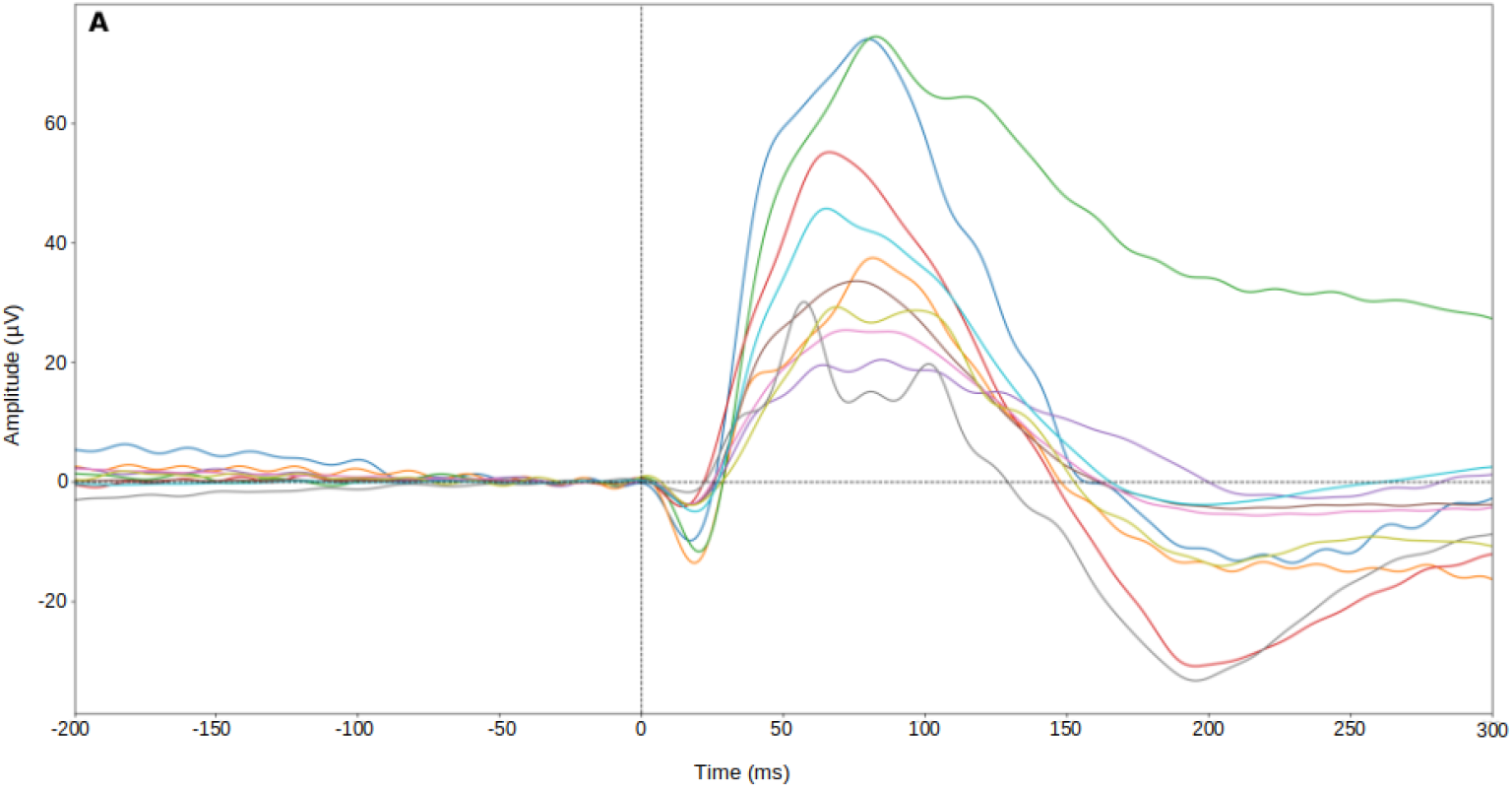
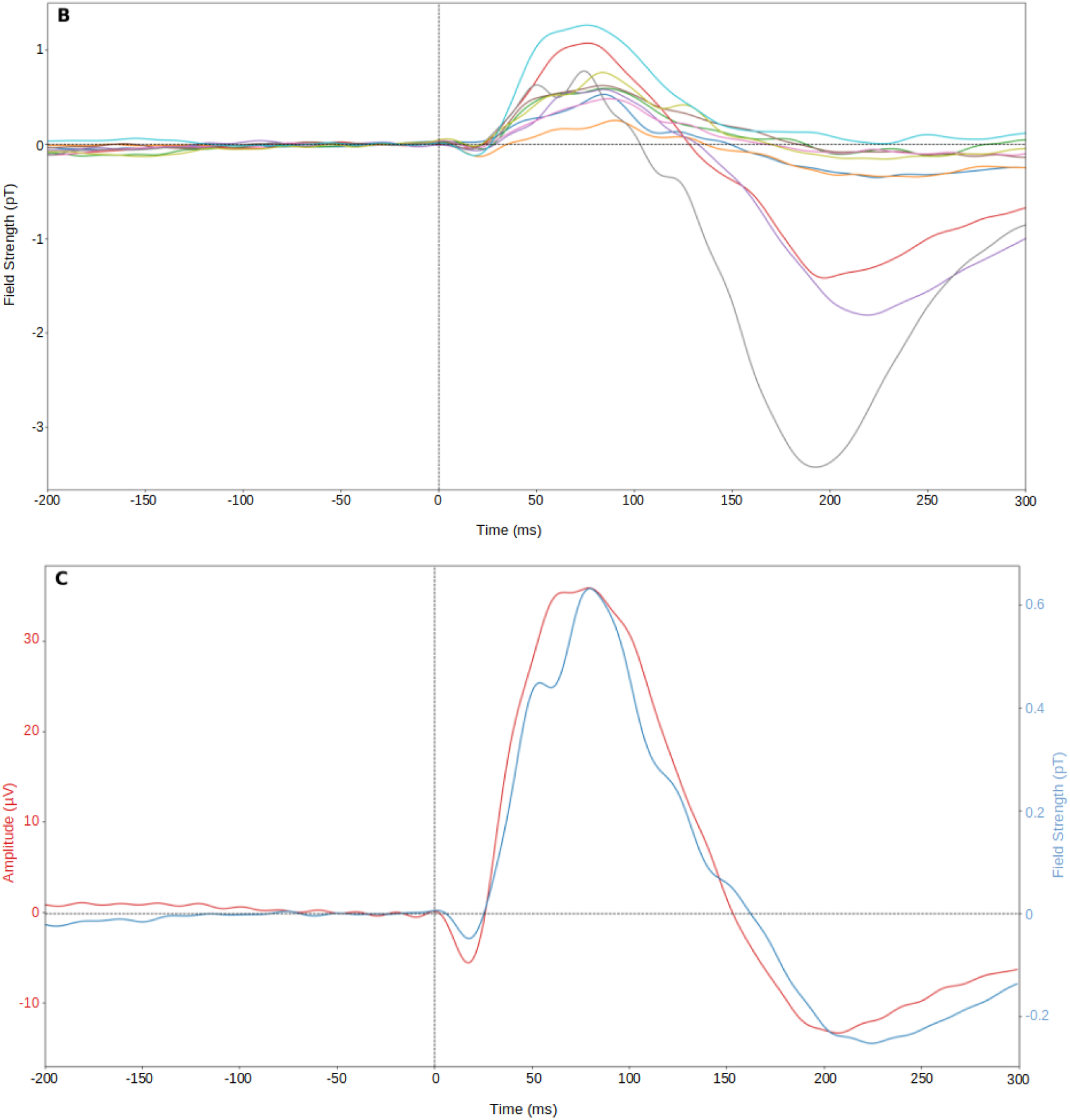
ERG and oMRG best sensors. **A**: Average of flash-evoked ERG per participant from under eye electrode **B**: Average of flash-evoked MRG per participant from the OPM with the highest amplitude responses across all participants (sensor 13). **C**: Grand average, ERG plotted in red and OPM sensor 13 in blue.

The robustness of the high frequency activity of the oMRG measurements was clearly maintained and is best seen in comparison to the ERG (Figure 3). Using the MRG sensor with the highest degree of similarity to the ERG, OPM 13, each participant’s data was filtered from 75 - 145 Hz (Figure 3A). The rHFO is visibly preserved in this channel with the oscillatory activity still visible even in the group average (Figure 3B). The ERG trace has been inverted to facilitate comparison with OPMs 1 and 4. In the MRG, the polarity of the signal is dependent on where the OPM sensor is relative to the retinal dipole (Armstrong & Janday, 1989). In our case, the OPM sensors above the eye showed a MRG trace with a negative a-wave and positive b-wave, with the opposite relationship for OPM sensors below the eye (data not shown).

**Figure 3.**
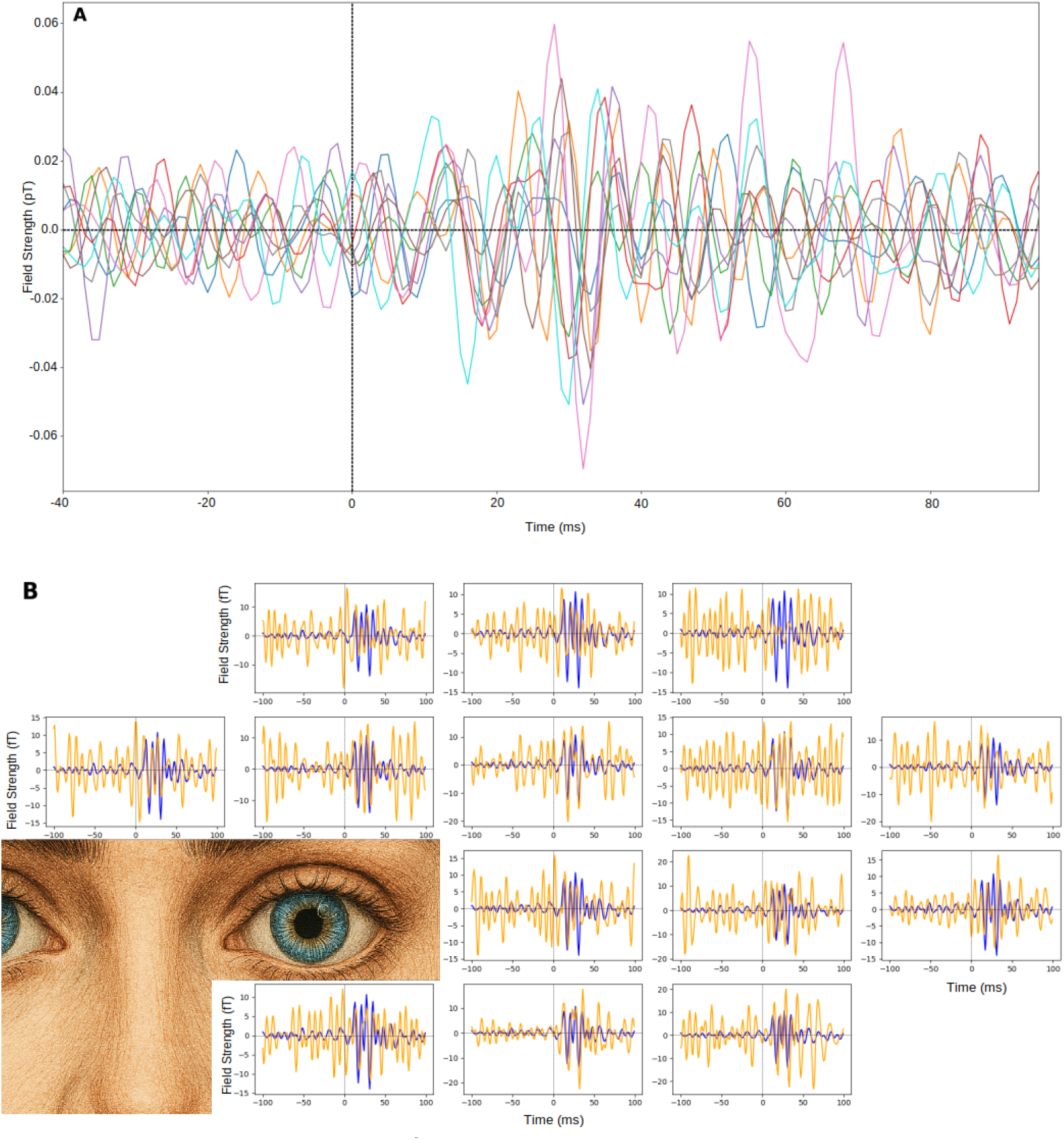
Evoked high frequency activity-. **A**: Individual traces from OPM 13, the OPM with the highest amplitude response filtered from 75 - 145 Hz. **B**: The grand average of each OPM (orange) and under eye ERG (blue), all signals were filtered from 110-145 Hz. Face only to aid with visualization, not to scale or anatomically accurate.

ITPC confirmed the oscillatory activity seen in the time series data was also present in the time-frequency domain. Figure 4 shows group ITPC results and the statistical effect of masking the group data with a critical p-value of 0.05. Sensors 5, 12, and 13 exhibited statistically significant responses following FDR correction within 50 ms of light flash onset.

**Figure 4.**
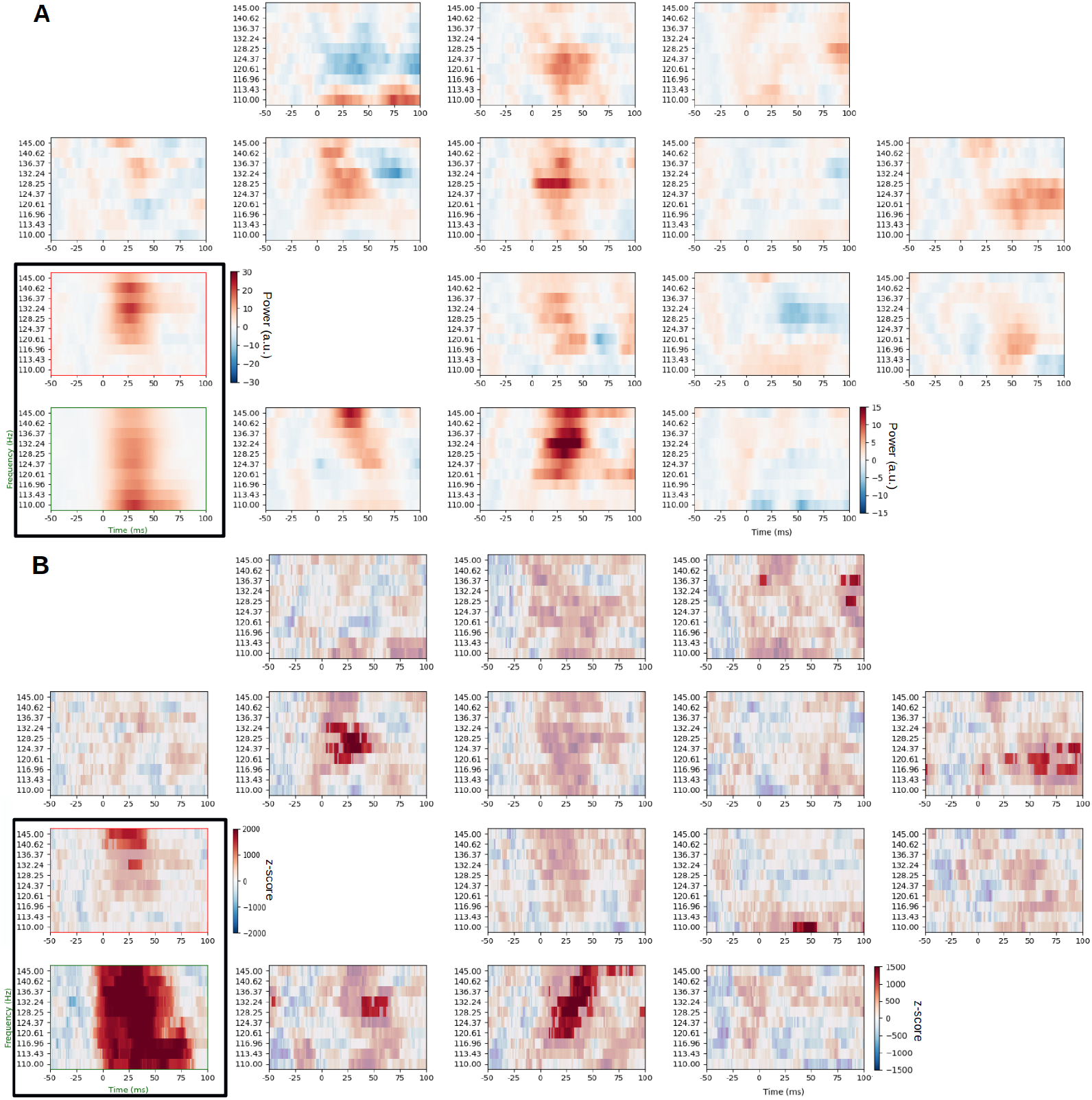
ITPC 110-145 Hz-. Above and below eye ERG plotted in black box in each subfigure; OPMs plotted as they were placed in array on face. The above eye ERG electrode border is in red and the below eye ERG electrode border is in green. The colorbar next to the above eye electrode (red plot border) applies for both ERG plots. The colorbar next to OPM 14 holds for all OPMs. **Top**: Grand Average ITPC plotted 110-145 Hz. **Bottom**: ITPC z-score plotted with insignificant areas masked, signed rank, alpha.05.

Sensor 5 was significantly active in the 124.37 Hz and 128.25 Hz bands, sensor 12 was significantly active in the 140.62 Hz band, and sensor 13 was significantly active in the 132.24 Hz, 136.37 Hz, 140.62 Hz, and 145.0 Hz bands (see figure 4).

Because significant activity in OPM 13 could possibly extend above 145 Hz, additional ITPC analyses above 145 Hz were run (Figure 5). First, 110 - 170 Hz was considered as this doubles the bandwidth being analyzed in the primary analysis from 35 Hz to 60 Hz. Figure 5a shows the statistical effect of masking the group data with a critical p-value of 0.05. Second, an ITPC analysis examining activity up to 300 Hz was run thus demonstrating that the extent of the analysis was accurate (Figure 5b).

**Figure 5.**
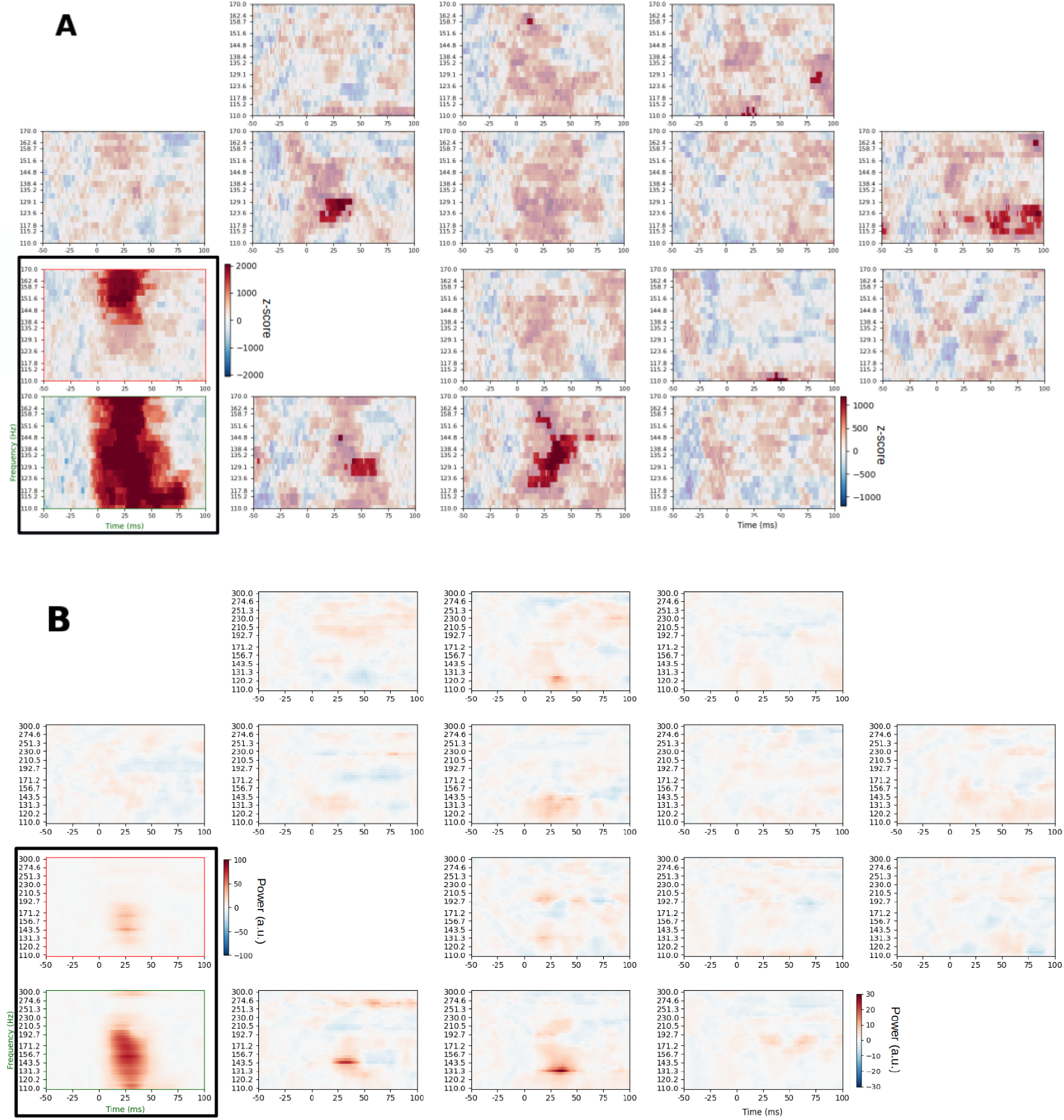
Activity above 145 Hz-. Above and below eye ERG plotted in black box in each subfigure; OPMs plotted as they were placed in array on face. The above eye ERG electrode border is in red and the below eye ERG electrode border is in green. The colorbar next to the above eye electrode (red plot border) applies for both ERG plots. The colorbar next to OPM 14 holds for all OPMs. **Top**: 110-170 Hz, ITPC z-score, 10 cycles, insignificant portions are masked, signed rank, and with an alpha of 0.05. **Bottom**: ITPC 110-300 Hz 15 cycles.

Analysis of the absolute amplitude spectrum, in OPM 13, demonstrated the frequency-dependent sensitivity of the OPMs, with detectable sensitivity extending up to approximately 165 Hz. This aligns with the observed significant oscillatory activity in the 120–158 Hz range of OPM 13 over the same time period (Figure 5a; grey shaded region in Figure 6). In this sensor, the amplitude follows a 1/f profile up to around 110 Hz, followed by an increase within the 120–160 Hz range, corresponding to the ITPC activity shown in Figure 5a. Comparison of the absolute amplitude spectra during the post-stimulus and pre-stimulus periods reveal that the OPM signal amplitude approaches the empirically estimated noise floor at approximately 165 Hz. This cutoff was calculated as where the lower standard error values of the post-stimulus spectra dropped below that of the pre-stimulus spectra for three continuous frequency bins (Figure 6; vertical dashed red line).

**Figure 6.**
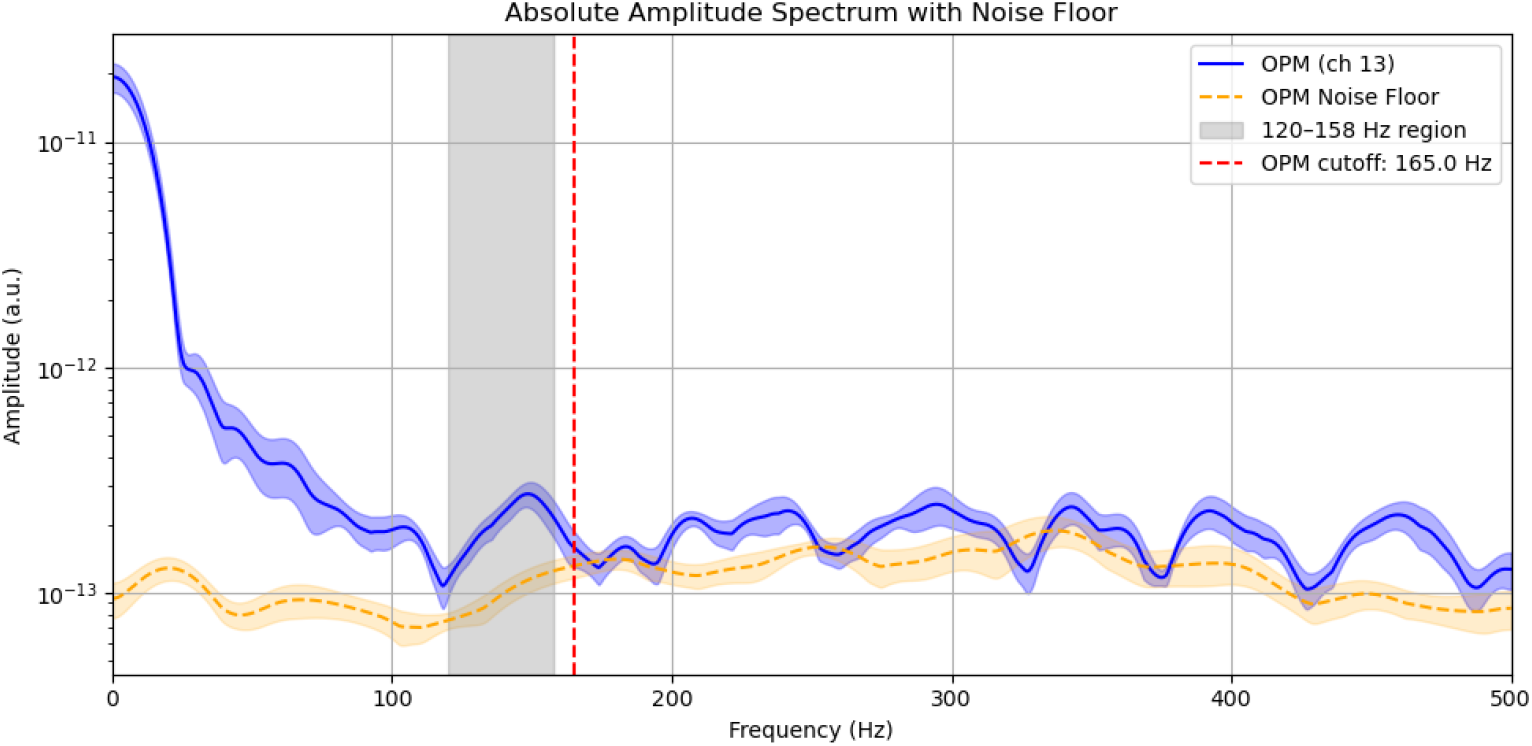
Absolute Amplitude Spectrum - In blue, the amplitude spectrum of channel 13 from 5 to 100 ms with standard error shading. In orange, the amplitude spectrum of the same channel computed from -50 to -5 ms; noise floor. The vertical red dashed line indicates the empirically determined OPM cutoff, where the lower standard error dropped below the noise floor for 3 consecutive frequency bins. The gray shaded region represents the frequencies in which the same OPM’s ITPC activity was significant during the same time window (see Figure 5a).

The mutual information between the under-eye and over-eye ERG electrodes was significant for all participants. MI between OPM sites and the under-eye ERG varied per participant and most OPM sites reflected shared information in the high-frequency bandwidth of interest 130 - 145 Hz. All participants exhibited significantly similar activity between the under-eye ERG and OPM 13 and OPM 14 (See supplementary material for more details). At the group level the grand average of the MI was computed and showed that 13 out of the 14 OPM sensors shared information significantly with the under-eye ERG (Figure 7). The insignificant OPMs were OPMs number 7 and 9.

**Figure 7.**
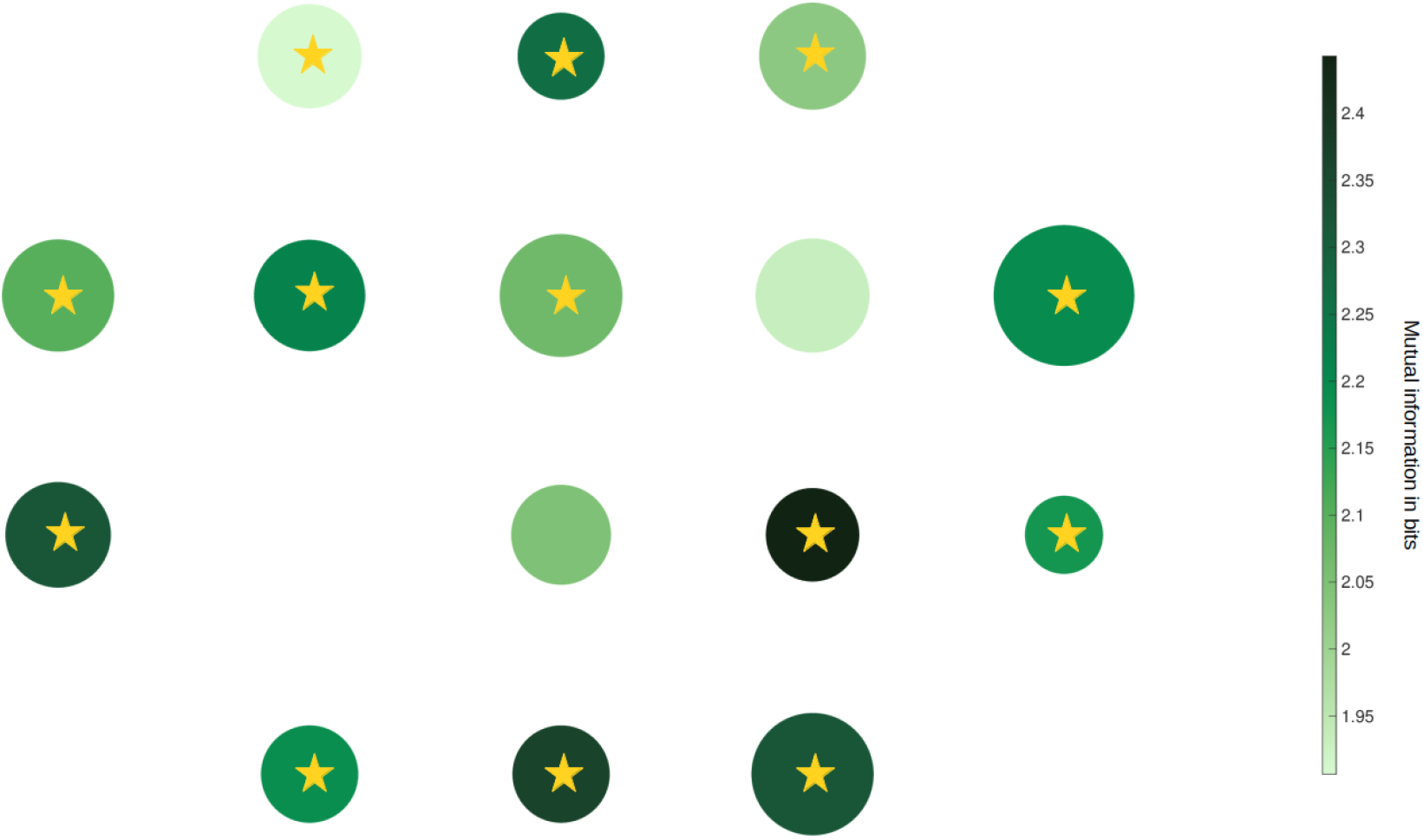
Mutual information per channel when compared with the under eye ERG activity-. Color corresponds to the mutual information in bits and the size of the circle is inversely proportional to the standard deviation across participants’ MI (i.e., smaller circles correspond to a less certain estimate of the mutual information). A significant amount of shared information is indicated by a gold star.

Additionally, MI between OPM locations and the under-eye ERG was compared using phase grand average ITPC data from the 110 - 145 Hz range. This analysis reveals the temporal and frequency characteristics of MI between the ERG and OPM sensors more clearly (Figure 8). There is strong alignment of information temporally following stimuli onset (1 ms) and again at around 50 ms. While temporally aligned, the band in which the activity varies depending on sensor location.

**Figure 8.**
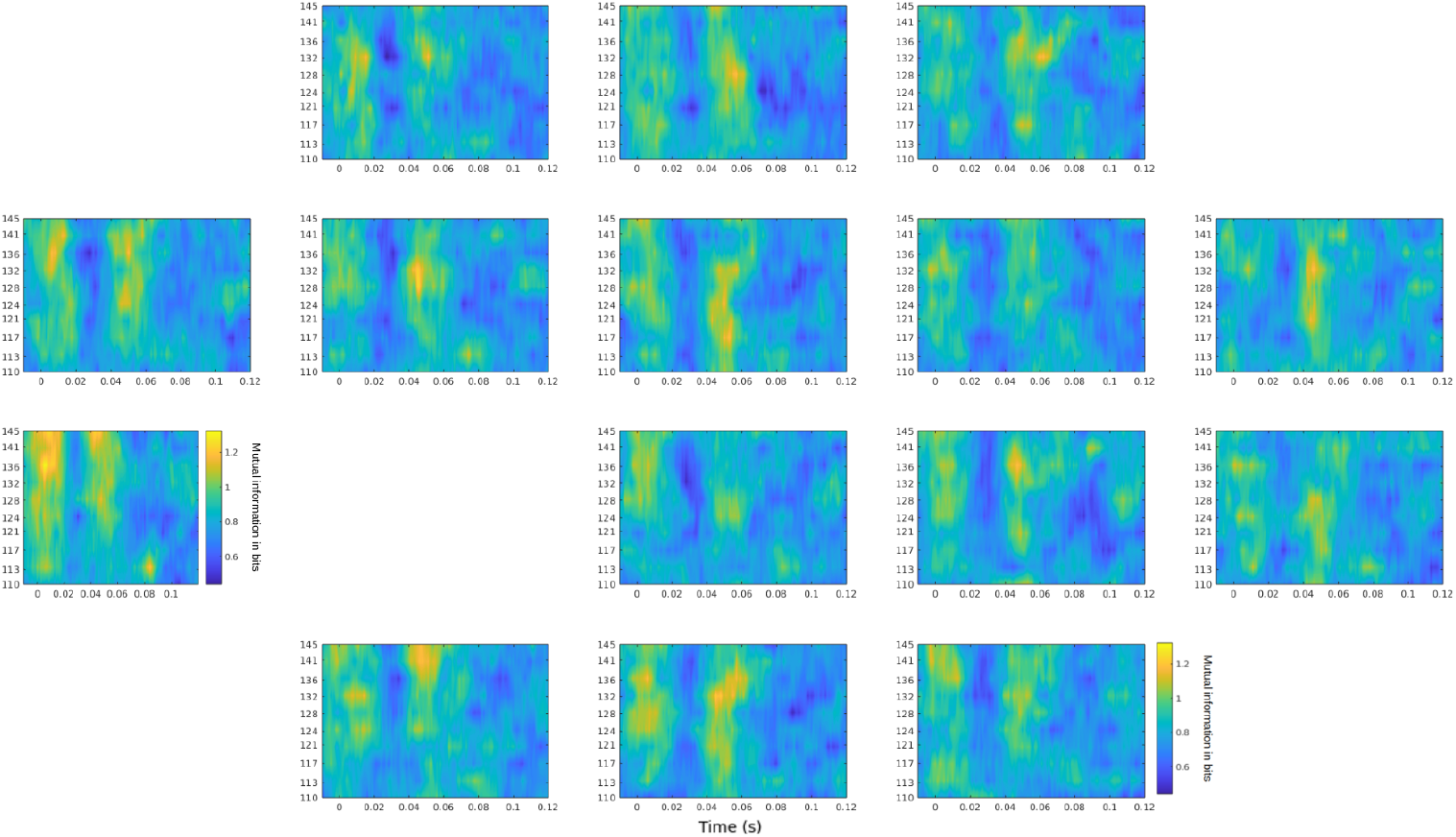
Mutual information of phase shared between under eye ERG and OPMs and above eye ERG from 110-145 Hz-. Window is constrained from -0.01 - 0.12 s and color bar is consistent for all plots.

## Discussion

In this study we measured high-frequency MRG responses from human participants to assess the sensitivity limits of modern OPMs and to validate that MRG with OPMs could meet clinical requirements. The retinal response to a flash of light produces a signal with well defined low and high frequency components (Robson et al., 2018). OPMs have already been shown to capture lower frequency components of MRG (Westner et al., 2021). In this study we measured evoked rHFO, the higher frequency component of MRG, as a litmus test for OPM sensitivity and we have shown that well placed OPMs can measure higher frequency biomagnetic signals. This also shows that OPMs can measure frequencies high enough to make the possibility of MRG a reality. By validating the MRG recordings with simultaneous ERG recordings we also provide a comparison. Across evoked representations of the signal, in the ITPC measurements, and in the mutual information measures there is agreement that MRG is sensitive to the same response as the ERG up to 145 Hz (Figures 3, 4, and 7). The upper limit of physiologically meaningful MRG signal detection appears to be around 158 Hz, as shown in Figure 5a, where the under-eye ERG electrode reveals broadband retinal activity and corresponding MRG responses remain observable up to this frequency. This is reinforced by the absolute amplitude spectra presented in Figure 6, which indicate that OPM sensitivity extends to 165 Hz. While there might seem to be an apparent discrepancy between the upper frequency bounds observed in the ITPC analysis (up to 158 Hz) and the absolute amplitude spectrum (up to 165 Hz), this likely reflects methodological differences. The ITPC analysis identifies statistically robust, phase-locked activity and was corrected for multiple comparisons, whereas the absolute amplitude spectrum reflects total signal power—including non-phase-locked components—and is not corrected for multiple comparisons. Therefore, while detectable signal energy may extend nominally up to 165 Hz, the physiologically meaningful, time-locked retinal response appears limited to ∼158 Hz. There also exists a significant patch of ITPC activity above 158 Hz (figure 5, OPM 2), within the relevant 0 - 750 ms time window, this activity is only seen in only one OPM that contains no other significant activity below 158 Hz and may very well be spurious (Figure 5a). As with most nascent technologies, this upper limit on OPM sensitivity is bound to improve (Gutteling et al., 2023).

In our measurements of the rHFO, we chose to utilize the OPMs’ closed-loop mode.

Activating closed-loop mode provides consistent -3 db frequency response over a wider bandwidth, a characteristic that measuring in an environment like the BMSR-2 can particularly benefit from. Reported sensor response for FieldLine V2 OPMs is -3 dB in open-loop mode and in closed-loop mode the usable bandwidth is defined by optimizing coefficients to best flatten the frequency response for each sensor (Robinson et al. 2022). In open-loop mode, rubidium based OPMs can be sensitive to fluctuations in the DC-150 Hz range. OPMs utilizing ^4^He atoms inside the vapor cell, operate in closed-loop and have demonstrated a wider available bandwidth (DC - 2000 Hz), but this is accompanied by trade-off in sensitivity and size (45 ft/√Hz; Gutteling et al., 2023). Theoretically, rubidium based OPMs operating in closed-loop mode can increase the sensitivity of an OPM to DC - 350 Hz, yet this is dependent on background fields that impact the range within which a sensor can functionally measure. Therefore, in environments less shielded than the BMSR-2, closed-loop mode could significantly degrade sensitivity. However, in the BMSR-2, which has a background gradient of 1.2 pT/mm and remaining field of less than 500 pT (https://www.ptb.de/cms/en/ptb/institutes-at-ptb/geraetezentrum-8-2.html), closed-loop mode should increase the measurable bandwidth without impacting sensitivity.

The benefits of performing measurements within the BMSR-2 represent another limitation, that is, without such a heavily shielded environment, the upper frequency components of MRG might be difficult to measure. Within a 3 layer shielded room (2x Mu-Metall shields and one Aluminum shield), like those typically used to record SQUID-MEG, lower frequencies (less than 50 Hz) and diagnostically valuable a- and b-wave components of the retinal response can be measured (Wester et al., 2021). Current work on improving the noise and gradient profiles of these modern 3 layer MSRs include new approaches to active and passive shielding that should create an environment suitable for high frequency measurements in the near future (Jas et al., 2025; Mellor et al., 2023; Seymour et al., 2022). Already, active shielding in combination with an OPM-optimized 3 layer MSR has created a one-cubic-meter recording volume zone with a remnant magnetic field of 670 ± 160 pT (Holmes et al., 2019; Rea et al., 2021). With shielding performance increases the likelihood of a fully developed MRG is possible.

Another aspect to consider is the angle that the OPMs were mounted on the participants (Figure 1). Both the over and under eye electrodes were mounted such that their sensing face pointed in towards the brain of the individual. The same was true for all OPM sensors. In our experiment, the sensing face of the OPM (the z-direction) was flush with the skin. While this is perhaps the easiest way to mount the sensors around the eye, the retinal dipole, which points the positive pole straight out of the pupil (Katila et al., 1981; van Schijndel et al., 1997) would be optimally measured if the primary sensing direction (B_z_) were positioned perpendicular to the dipole. Many OPM designs now allow measurements along all three axes and future MRG work will benefit from the ability to measure from tangential and radial sources (Boto et al., 2022; Feys et al., 2023; Gutteling et al., 2023; Rea et al., 2022)).

Here we have detailed the bandwidth of zero-field OPMs in closed-loop mode for measuring MRG. Our results conservatively demonstrate that current zero-field OPM technology can resolve neural activity up to ∼145 Hz, supporting their use not only for MRG, but also for MEG measurements of cortical high gamma activity. Given that OPM-based MRG measures similar features as ERG, including high-frequency oscillations, it can serve as a comfortable alternative for clinical and neuroscientific examinations. It may also simplify combination of retinal and cortical measurements for investigating information transfer across the early visual pathway (Westner et al., 2019). Finally, the topographic information gathered by an OPM array around the eye promises to yield spatial detail beyond ERG, with the potential for delineating the retinal neurons involved in generating high-frequency oscillations.

## Acknowledgements

This work was supported by an ERC Starting Grant (ERC-StG-640448) to SSD and a Danish Diabetes Foundation Travel Grant awarded to JIL. The Core Facility ‘Metrology of Ultra-Low Magnetic Fields’ at Physikalisch-Technische Bundesanstalt receives funding from the Deutsche Forschungsgemeinschaft – DFG (funding codes: DFG KO 5321/3-1 and TR 408/11-1)

## Ethics

This study was conducted at Physikalisch-Technische Bundesanstalt in Berlin, Germany. Protocol was approved by the PTB Ethics Committee application number 2019-1.

## Code and Data Availability

Anonymized data is available on request. Code can be found at https://github.com/Lubell/rHFO_OPMs

## Author Contributions

JIL conceived, designed, and executed the experiment, analyzed data, and wrote the manuscript. JV and KR were involved with data collection and recruitment. TS was involved with the design of the experiment and data collection. SSD conceived of the study and was involved with experimental design and wrote the manuscript.

## Supplementary Material

Here we only report the non-significant results for mutual information (see figure 7 and table 1 below).

**Table 1.**

| Participant number | Non-significant OPM sites |
| --- | --- |
| 1 | 2, 7 |
| 2 | 1, 10, 12 |
| 3 | none |
| 4 | 2, 12 |
| 5 | none |
| 6 | 2, 3, 4, 5, 9, 11 |
| 7 | none |
| 8 | 8 |
| 9 | 4, 11 |

**Supplementary Figure 1:**
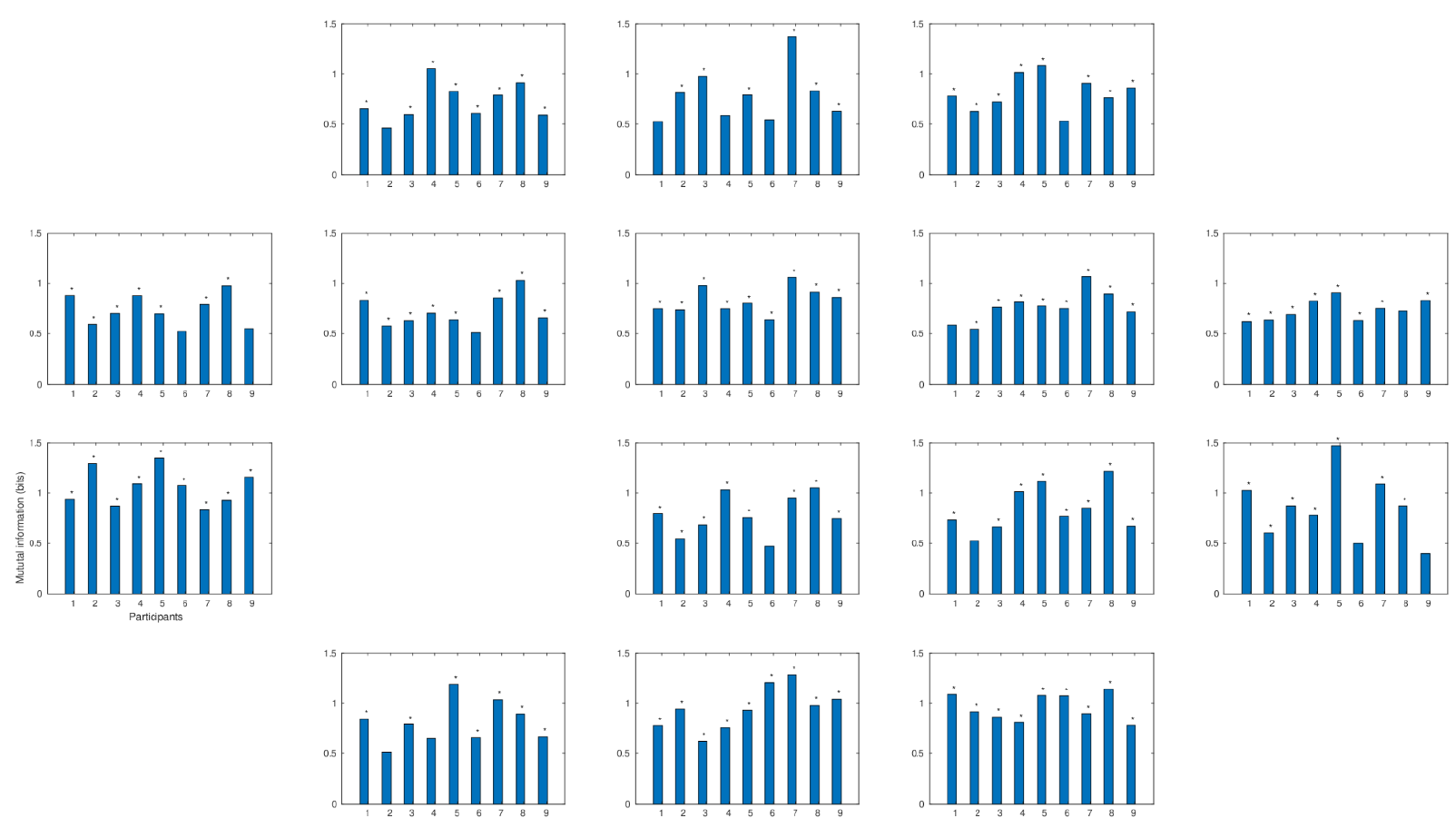
Amount of mutual information in bits between the under-eye ERG electrode and each sensor (including over eye electrode) for each participant, significant sensors marked with ‘*’ less than.0025. Significance determined using permutation testing.

